# Mycoplasma stress response: adaptive regulation or broken brakes?

**DOI:** 10.1101/004960

**Authors:** Pavel V Mazin, Gleb Y Fisunov, Alexey Y Gorbachev, Ilya A Altukhov, Tatiana A Semashko, Dmitry G Alexeev, Vadim M Govorun

## Abstract

The avian bacterial pathogen *Mycoplasma gallisepticum* is a good model for transcriptional regulation studies due to its small genome and relative simplicity. In this study, we used RNA-Seq experiments combined with MS-based proteomics to accurately map coding sequences (CDSs), transcription start sites (TSSs) and transcription terminators (TTs) and to decipher their roles in stress-induced transcriptional responses. We identified 1061 TSSs at an FDR (false discovery rate) of 10% and showed that almost all transcription in *M. gallisepticum* is initiated from classic TATAAT promoters, which are surrounded by A/T-rich sequences and rarely accompanied by a −35 element. Our analysis revealed the pronounced complexity of the operon structure: on average, each coding operon has one internal TSS and TT in addition to the primary ones. Our new transcriptomic approach based on the intervals between the two closest transcription initiators and/or terminators allowed us to identify two classes of TTs: strong, unregulated and hairpin-containing TTs and weak, heat shock-regulated and hairpinless TTs. Comparing the gene expression levels under different conditions (such as heat, osmotic and peroxide stresses) revealed widespread and divergent transcription regulation in *M. gallisepticum*. Modeling suggested that the structure of the core promoter plays a major role in gene expression regulation. We have shown that the heat stress activation of cryptic promoters combined with the suppression of hairpinless TTs leads to widespread, seemingly non-functional transcription.

## Introduction

*Mycoplasma gallisepticum* belongs to the Mollicutes class – a specialized branch of microorganisms related to Gram-positive bacteria (Davis et al. 2013). *M. gallisepticum* is an important pathogen in poultry and wild birds, in which it causes chronic respiratory disease (Levisohn and Kleven 2000). Mollicutes feature reduced genomes with an average size of 1 Mb, and they lack a cell wall (Razin and Yogev 1998). Consequently, their cell physiology is considerably simplified compared with that of most bacteria, making Mollicutes a good model for systemic studies and, in particular, for studying the complex response to stress.

*M. gallisepticum*, along with most Mollicutes, shows a reduced repertoire of transcription factors (TFs) compared with that of related bacteria, such as *B. subtilis* (Moreno-Campuzano et al. 2006). The only TF whose mechanism is known in *M. gallisepticum* is a heat-shock repressor (HrcA) that binds a palindromic sequence (CIRCE) in the promoters of several chaperone genes (Chang et al. 2008), whereas other common bacterial TFs, such as the LexA repressor of the SOS response (Carvalho et al. 2005), are lacking. Recent studies that have demonstrated widespread differential expression in response to a variety of stresses in the *Mycoplasma* species (Güell et al. 2009; Weiner III et al. 2003; Gorbachev et al. 2013) raise questions about the underlying regulation of these responses. Experimental identification of transcription start sites (TSSs) and transcription terminators (TTs) could help to resolve this “regulation without regulators” puzzle.

Classic views on bacterial transcription assume that coding sequences are organized into operons – genomic regions that are transcribed as a single RNA (Jacob and Monod 1961). On the contrary, recent genome-wide transcriptomics studies on several bacterial species (*E. coli* (Cho et al. 2013), *B. subtilis* (Kobayashi et al. 2007), *Listeria monocytogenes* (Toledo-Arana et al. 2009)*, Helicobacter pylori* (Sharma et al. 2010) and *Mycoplasma pneumoniae* (Güell et al. 2009)) demonstrated that operons frequently include internal promoters and TTs that lead to the so-called “staircase” transcription (Güell et al. 2009). These results cast doubts on the operon paradigm and call for a new, more realistic approach.

Common RNA-Seq experiments do not allow for the precise identification of TSSs; however, several techniques that address this problem have recently been developed. These techniques include either tagging of the 5′-ends by the ligation of a specific adapter (Cho et al. 2013) or 5′-end enrichment procedures using a 5′-phosphate-dependent nuclease (Sharma et al. 2010).

In the current work, we systematically investigated transcription and translation in *M. gallisepticum* under several conditions using common RNA-Seq experiments, 5′-end enriched RNA-Seq and mass spectrometry-based proteomics. We introduced a new approach to study bacterial transcription that is based on *intervals* (genomic regions between the two nearest TSSs and/or TTs) instead of operons. By combining both experimental evidence and *in silico* prediction, we identified hundreds of promoters, ribosome-binding sites (RBS), TTs, operons and non-coding RNAs (Fig. 1A).

**Figure 1.**
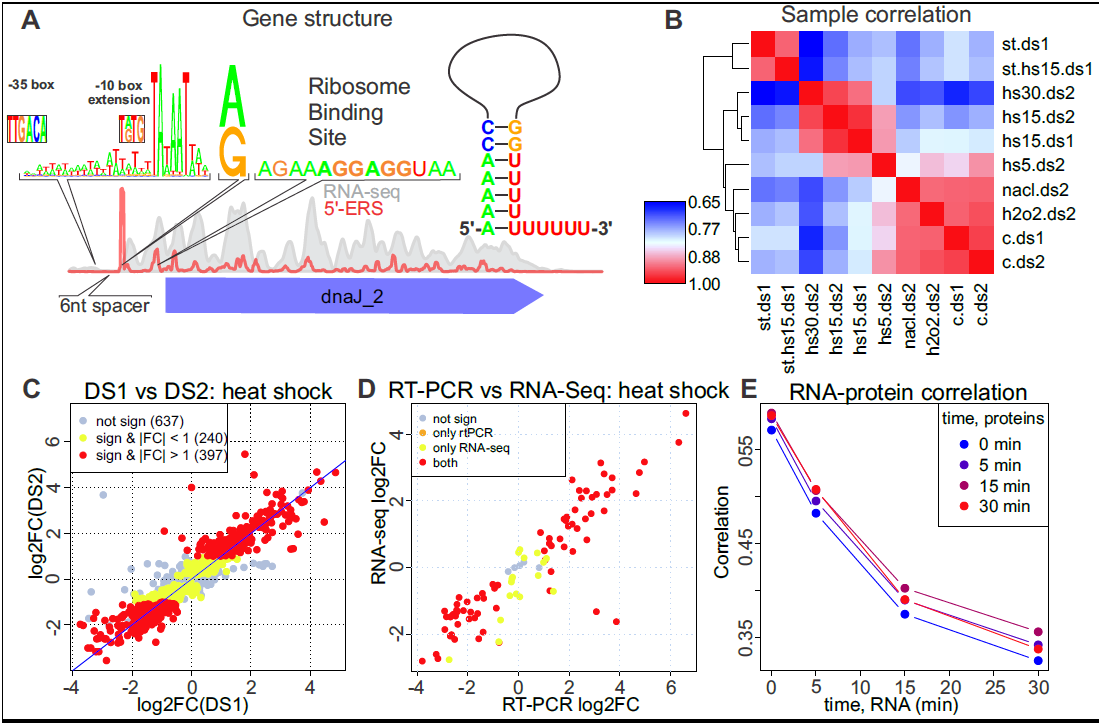
(A) Typical RNA-Seq and 5’-ERS coverage profiles near the dnaJ2 gene. The diagram of the promoter sequence, the first nucleotide of the transcript, RBS and a schematic representation of the TT are shown above the plot. (B) Correlation heatmap (correlation increases from 0.65 (blue) to 1 (red); one minus the Spearman correlation coefficient was used as the distance metric for clustering) for different conditions (c, h2o2, nacl, hs5, hs15 and hs30 denote the control, oxidative stress, osmotic stress and heat stress for 5, 15 and 30 min, respectively) and datasets (ds1 and ds2). The expression values represent the average values of the replicates. (C) Agreement of the heat-shock (15 min)-related expression changes (log2-fold change) between DS1 (x-axis) and DS2 (y-axis). One dot denotes one CDS. Not significant, significant but with a fold change below 2 and significant with a fold change above 2 are shown in gray, yellow and red, respectively. (D) Agreement of the heat-shock (15 min)-related expression changes between RT-PCR (x-axis, differences in cycle number) and DS2 (y-axis, log2-fold change). The same color scheme as in panel C was used. (E) Dependence of the Spearman correlation coefficient between the protein abundance (PAI) measured for different heat shock durations (shown with different colors) and the mRNA abundance (RPKM) on the heat shock duration used for the RNA-Seq experiments.

## Results

In this study, we strand-specifically sequenced two batches of samples (16 samples in each batch) of the total RNA extracted from *M. gallisepticum* that was cultured under different conditions (exponential and stationary growth phases; heat shock, osmotic and oxidative stresses) in at least two biological replicates using a SOLiD 4 sequencer (Life Technologies). The sequencing produced 989 million reads, 44% of which could be mapped to the genome. Although most of these reads correspond to rRNA and tRNA, 17% are mapped to CDSs and 2.6% are mapped to intergenic regions (Supplementary Table S1). In addition to common RNA-Seq experiments, we employed a 5’-end enrichment procedure (see Methods) to precisely identify the 5′-ends of the transcripts (5’-enriched RNA sequencing, 5’-ERS). Correlation analysis shows good sample clustering ruled by the biological conditions rather than by technical variations (Fig. 1B, Supplementary Fig. S1). Our analysis shows a close agreement between the gene expression changes under heat shock measured in the two datasets (the Spearman correlation coefficient between the log fold changes is 0.92; Fig. 1C). Additionally, we performed RT-PCR for 98 selected genes under the same conditions (Supplementary Table S2). The gene expression levels as well as the fold changes measured by these two techniques show high correlation (rho > 0.7, Fig. 1D, Supplementary Figs. S2 and S3). We used mass-spectrometry (see below) to assess the protein abundance index (PAI) under the heat shock and control conditions. The correlation between the PAI and mRNA abundance (RPKM) is approximately 0.5 for the control conditions (Fig. 1E), which is comparable to the estimates obtained in other works (Maier et al. 2011). The foregoing results confirm the high quality of our data and support their application in characterizing the transcription organization in *M. gallisepticum* strain *S6*.

### Operon organization in *M. gallisepticum*

Traditionally, bacterial transcription is considered to be organized into operons, which are genomic regions with a single promoter and several coding sequences that are transcribed as a single mRNA (Jacob and Monod 1961). In fact, bacterial transcripts seem to overlap with each other (Toledo-Arana et al. 2009; Cho et al. 2013), resulting in a complex structure with internal TSSs and/or TTs. Here, we define operons as a set of overlapping transcripts. Operons can be divided into the intervals between two nearest TSSs or TTs. Theoretically, the read coverage in such intervals should follow a Poisson distribution, but practically, their distribution is much more variable due to the differences in nucleotide composition, RNA secondary structure and sequencing biases (Khrameeva and Gelfand 2012). We used generalized linear models (GLMs) with a quasi-Poisson distribution to model the read coverage. We applied a quasi-log likelihood test to divide the *M. gallisepticum* transcriptome into 1059 equally covered intervals (see Methods, Fig. 2A and Supplementary Table S3). The borders between the intervals are formed by either up- or down-coverage steps (up-CSs or down-CSs), which should theoretically correspond to TSSs and TTs, respectively. In total, we observed 499 up-CSs and 558 down-CSs. We assigned annotated features (CDS, tRNA and rRNA) to the intervals if the overlap was greater than 50% of the length of the gene. Of 877 annotated genes, 868 could be assigned to intervals and 586 reside completely within a single interval. The read coverage strikingly differed between the intervals that contained genes and those that did not (Fig. 2B). Assuming that 95% of the gene-containing intervals are expressed, we set the coverage threshold for expressed intervals to 2.38 read per position. The 372 intervals with a coverage value below the threshold were considered to be unexpressed. Most of the intervals overlap with just a few genes. The longest (in terms of the number of overlapped genes) expressed interval (12,638 nt) contains 24 genes that encode ribosomal proteins (Fig. 2C). The definition of an operon as a continuous sequence of expressed intervals results in 208 operons, 125 of which overlap 839 genes (in total) by at least 50% of the gene length. Furthermore, 772 genes reside completely within an operon. In addition to the 208 primary up-CSs and down-CSs, the operons contain 218 and 261 internal up-CSs and down-CSs, respectively. Most of these CSs are observed within gene-containing operons, most likely because the latter are usually much longer than the operons that do not contain genes (Fig. 2D). We compared our prediction for each consecutive pair of genes that resided within the same or in different intervals with the operon prediction obtained from proOpDB (Taboada et al. 2012) and observed a high and significant overlap (Fisher exact test p < 1e-14). The excess of positive correlation values for the gene expressions across the different conditions for the CDSs from the same interval (and, to some degree, from the same operon) compared with the correlation for the genes from different operons (Fig. 2E) confirms the robustness of our procedure.

**Figure 2.**
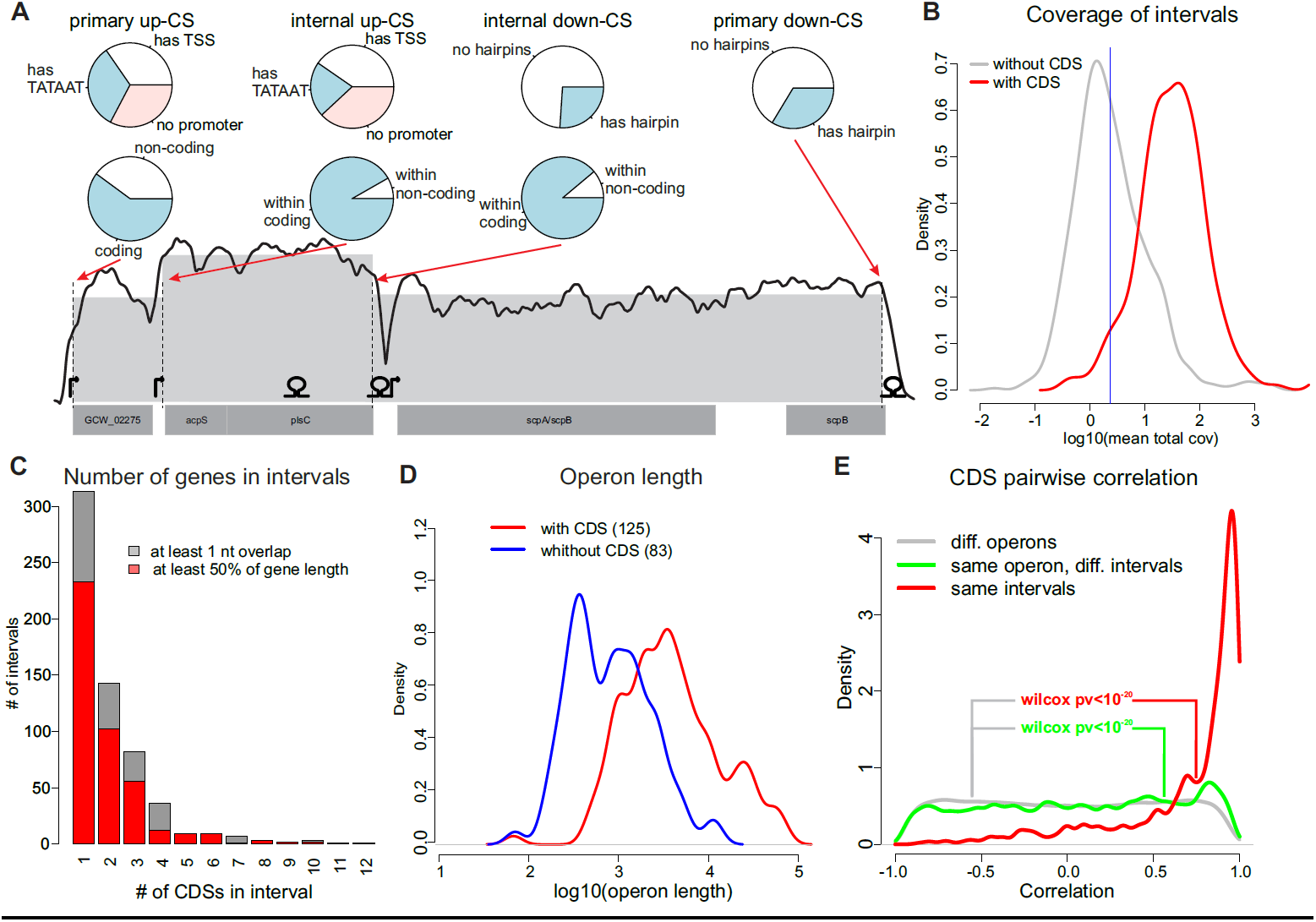
Operon prediction. A) Example of a predicted operon. Annotated CDSs are shown at the bottom; the TSSs identified by 5’-ERS and the hairpins predicted by RNIE are shown above. The smoothed coverage (running mean in a 100 nt window; log scale) is shown with a solid line; the dashed vertical lines represent up- and down-CSs; and the mean interval coverage is represented by the gray area. Classifications of primary and internal CSs by TSS/hairpin existence and the coding potential of the operons to which they belong are shown with pie charts. B) Distribution of intervals with (red) and without (gray) CDSs by log coverage. The 5% quantile of the former is shown as a blue vertical line. C) Distribution of intervals by the number of genes. D) Distribution of operons with (red) and without (blue) CDSs by log length. E) Distribution of Pearson correlation coefficients for the pairs of genes that belong to the same interval (red), same operon (but not interval, green) and genes from different operons (gray).

### Promoter structure in *M. gallisepticum*

To further understand the transcription organization in *M. gallisepticum*, we used our 5’-ERS samples to identify the exact locations of TSSs. Briefly, we looked for the local maxima (peaks) in the 5’-ERS coverage, optimized the position weight matrix (PWM) for the upstream region and then used the PWM to identify the peaks with a significantly better PWM match than expected by chance (see Methods). The procedure resulted in 1061 TSSs at an FDR < 10% (Supplementary Table S4). All of the identified promoters are sigma 70 promoters with a TA[T/A]AAT −10 element surrounded by an A/T-rich region. We searched for the TRTGN 5’-extension of the Pribnow box that was previously reported for *B. subtilis* and other Gram-positive bacteria (Voskuil and Chambliss 2002). We found 25 TSSs (out of 1061) with such an extension; this result is 5-fold greater than expected by chance (see Methods). The primary sigma factor of *M. gallisepticum* (RpoD) along with the Pribnow box-binding domain contains the −35 element-binding domain (Supplementary Fig. S4). To test whether *M. gallisepticum* promoters contain the −35 element, we searched up to 100 nt upstream of TSSs for a consensus TTGACA sequence (Hinton 2007). The results indicated that 122 TSSs (2-fold more than expected by chance) have a TTGACA sequence with no more than two mismatches at the −35 position. An analysis of the *M. gallisepticum* genome revealed one putative sigma factor-like transcription factor (GCW_00440) that may function as an alternative sigma factor. However, the regions around the 5’-ERS peaks did not exhibit the enrichment of motifs that were dissimilar to TATAAT, leading us to the conclusion that *M. gallisepticum* has only one functional sigma factor and that the observed TATAAT-lacking 5’-ERS peaks represent experimental noise. An analysis of the promoter structure revealed that the −10 element is separated by a 6 nt (or, rarely, a 7 or 5 nt) spacer from the TSS (see Methods, Fig. 3A and Supplementary Fig. S5). Similarly to transcription in *Mycobacterium tuberculosis* (Cortes et al. 2013), transcription in *M. gallisepticum* is initiated solely from an A or G nucleotide. If the 7th position from the −10 element is occupied by a C or T, transcription initiates one nucleotide downstream, resulting in a 7 nt spacer. Unfortunately, there is no such simple explanation for the most rare spacer, the 5 nt spacer (Fig. 3A). The promoters predicted based on 5’-ERS are located significantly closer to the up-CSs than the same promoters that are randomly shifted by no more than 200 nt (Wilcoxon test, p < 1e-8, Fig. 3B). Although only 35% of the up-CSs have promoters that are predicted from 5’-ERS within a ± 20 nt interval, most of them (62%) have a good PWM match in the same region. Interestingly, most of the TSSs identified by 5’-ERS (857 TSSs, 81%) are not associated with up-CSs. Although some of these 857 TSSs might be false positives, 69% of them are associated with the increase in read coverage and are likely to contribute to transcription. The 5’-ERS-identified TSSs that are not associated with up-CSs have a low relative step size and coverage by 5’-ERS reads (six-fold lower on the average than the coverage of the up-CSs associated TSSs) and constitute only 43% of the overall transcription initiation activity. Similarly, the up-CSs that lack 5’-URS-identified TSSs have a lower step size than the up-CSs that have a TSS within 20 nt (Fig. 3C).

**Figure 3.**
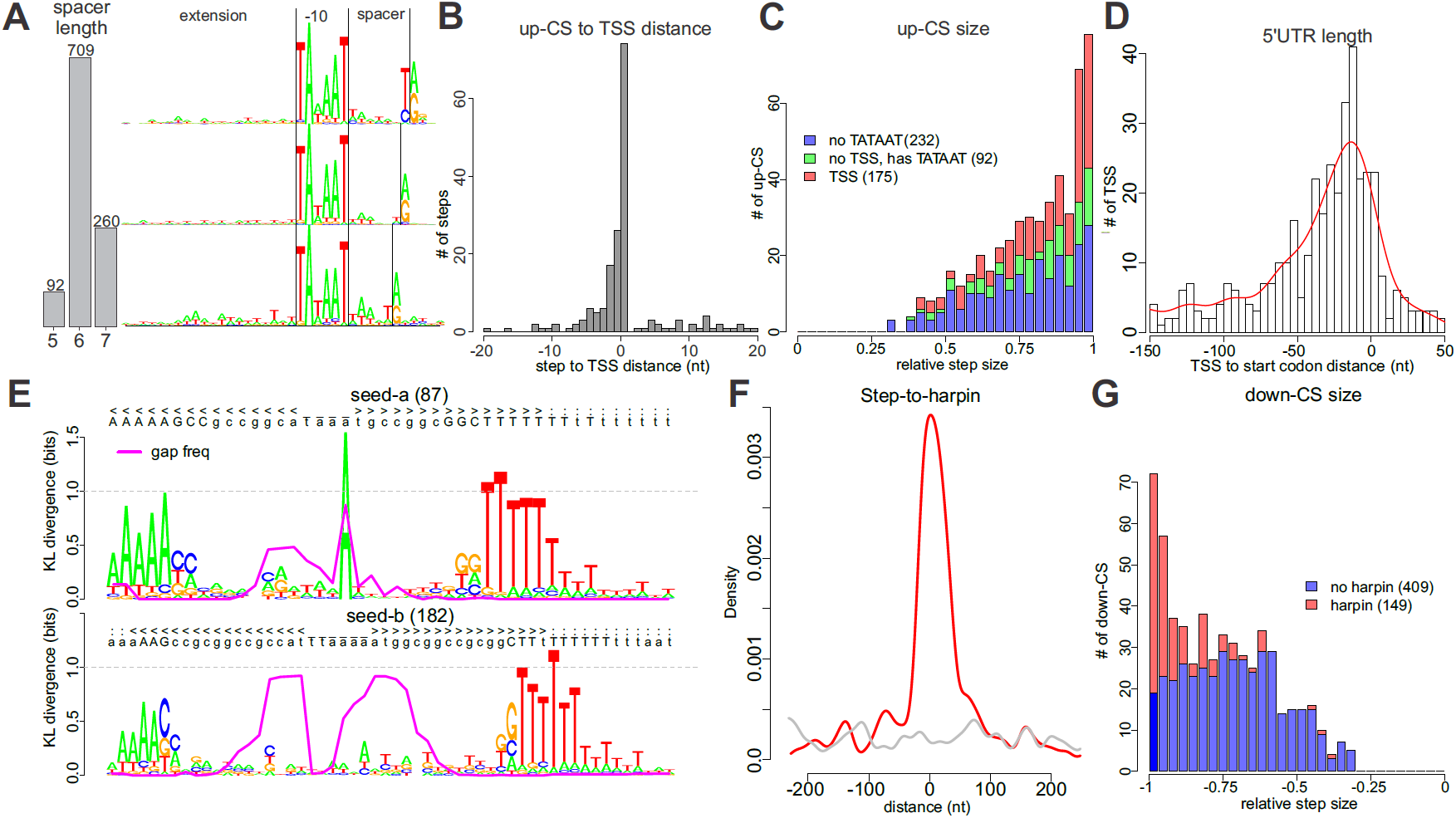
Structure of *M. gallisepticum* TSSs and TTs. A) Distribution of TSSs by spacer (between the −10 element and the TSS) length (left) and logo-images of the promoter region for each spacer length (7 nt to 5 nt from top to bottom). The −10 element and TSS are shown with vertical black lines. B) Distribution of distances between the TSSs and the nearest up-CS. C) Distribution of up-CSs by step size. The up-CSs that have 5’-ERS-detected TSSs, have only a good −10 element (detected by PWM) and have no signs of a TATAAT-like promoter are shown in red, green and blue, respectively. D) Distribution of distances from the nearest start codon to a TSS (negative values correspond to TSSs placed before the start codon). E) The logo-images of alignments of RNIE-predicted TTs to two seeds. The proportion of gaps in a given position is shown with a magenta line. F) Distribution of distances between down-CSs and RNIE-predicted hairpins. G) Distribution of down-CSs by relative step size. The down-CSs with and without hairpins are shown in red and blue, respectively.

We observed that 39% of the predicted TSSs reside in close proximity (<200 nt) to a start codon. The distribution of the 5’ UTR length has a mode near 15 nt, and nine transcripts appear to be leaderless (Fig. 3D). For eight of these transcripts, we detected peptides from the corresponding proteins in the proteomic data; three of these proteins are the first ORFs in polycistronic transcripts.

### Structure of transcription terminators in *M. gallisepticum*

Because no ORF-encoding Rho factor was found in the *M. gallisepticum* genome, the transcription termination in *M. gallisepticum* should occur in a Rho-independent manner and is likely to be associated with RNA secondary structures, such as hairpins (intrinsic terminators) (Farnham and Platt 1981). We predicted 256 hairpins genome-wide using the RNIE program (Gardner et al. 2011). Our results suggest that the stems of the terminator hairpins in *M. gallisepticum* are reduced compared to those of the default RNIE models and usually consist of 5 A-T pairs and just two G-C pairs near the loop (Fig. 3E).

The predicted hairpins cluster near down-CSs (Fig. 3F) as well as near the stop codons. We observed a weak but significant (rho=-0.18, p < 0.002) correlation between the hairpin score and the relative coverage step size. The hairpins that cannot be associated with a down-CS have a significantly lower score (one-sided Wilcoxon test, p < 0.03) than that of the hairpins that reside within a ± 200 nt interval from a down-CS. Interestingly, whereas most of the down-CSs with a relative step size close to −1 have a hairpin (102 of 166 down-CSs with a step size below −0.9), a large class of down-CSs with a step size distributed around −0.7 mostly lacks hairpins (only 47 of 392 down-CSs with a step size above −0.9 have a hairpin; Fig. 3G). Both classes of down-CSs are accompanied by a sufficient decrease in the GC content (from 32% in the upstream region to 27% in the downstream region; Supplementary Fig. S6).

### Transcriptional response of *M. gallisepticum* to stress

We used annotated CDSs to assess the differential expression in *M. gallisepticum*. All normalizations used in RNA-Seq-based transcriptomics assume that only a tiny fraction of the genes change their expression or, at least, that the numbers of up- and down-regulated genes are approximately equal (Dillies et al. 2013). Here, we used RLE normalization from the edgeR package (Robinson et al. 2010) to scale the library sizes. When applied to all samples but the stationary phase, the calculated normalization factors are within the 0.82-1.2 interval, and their effect is below our fold change threshold (2). However, if the stationary phase samples are included, the results change dramatically: the normalization factors for the stationary phase samples (0.13-0.32) are approximately ten times lower than the normalization factors for other samples (1.9-2.7). Such huge differences may arise if most of the genes are down-regulated in the stationary phase. Our RT-PCR experiments confirm this idea: when compared to 23S rRNA abundances, the abundances of most mRNAs decrease in the stationary phase (Supplementary Fig. S3). Because no normalization could be applied in such circumstances, we did not scale the library sizes for differential expression analysis under the stationary phase (see Methods). We used the SAJR package (Mazin et al. 2013) to perform a pairwise comparison for the control samples and each of the stresses. Genes with a q-value (BH-corrected p-value) below 0.05 and fold change above two were considered differentially expressed (for both up- and down-regulation).

The most significant change in the transcriptional landscape of *M. gallisepticum* was observed upon the transition to the stationary phase (Fig. 4A, Supplementary Table S5). For the stationary phase, we identified 723 down- and 30 up-regulated genes. The up-regulated genes were significantly enriched in genes associated with the “response to stress,” “oxidation-reduction process” and “glycolysis” (Supplementary Table S6). Only 67 and 118 genes change their expression under oxidative and osmotic stresses, respectively, and most of these genes are up-regulated (40 and 77, respectively).

**Figure 4.**
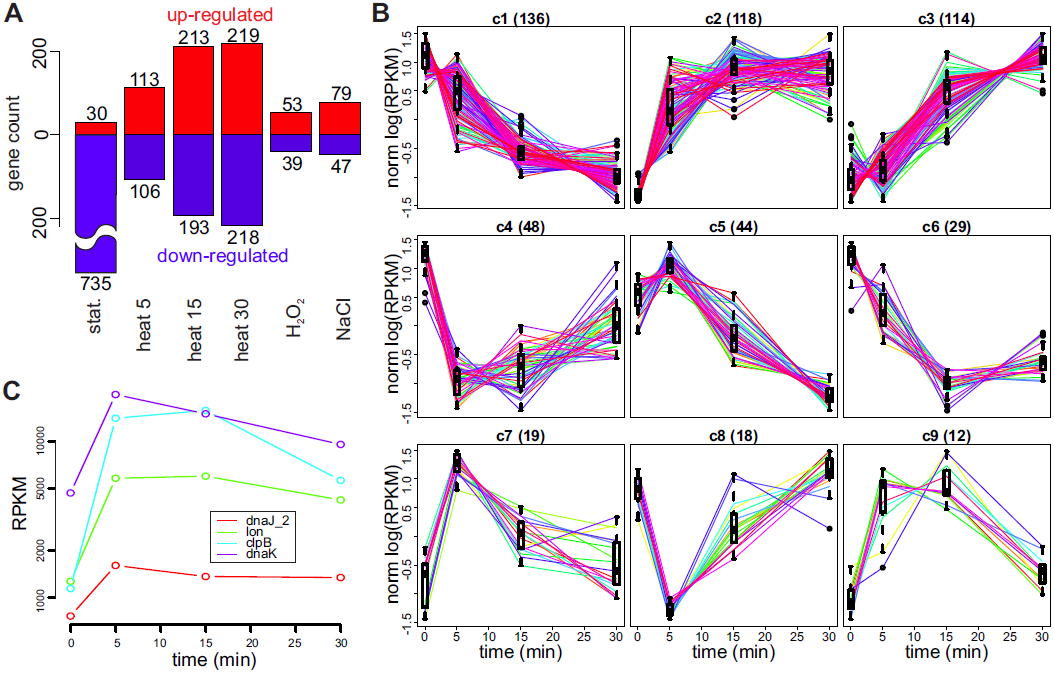
Transcription regulation. A) Number of CDSs whose expression changes significantly under different conditions. The up- and down-regulated genes are shown in red and blue, respectively. B) Nine patterns of gene expression changes under heat shock. The patterns are ordered by their size (shown in brackets in the panel titles). The normalized log expression of individual genes is shown with lines, and the distribution of the normalized log expression at each time point is shown with a box. C) Gene expression profiles (RPKM) of four genes that have the CIRCE motif in the upstream region.

Genes that are up-regulated under oxidative stress are significantly enriched in the “iron-sulfur cluster assembly” and “oxidation-reduction process” GO terms. These genes encode the Fe-S cluster assembly protein (SufB), the scaffold protein for Fe-S clusters assembly (NifU), methionine-sulfoxide reductase (MsrB), azoreductase (AsoR), cysteine desulfurase (CsdB), flavodoxin and several other proteins involved in ROS protection (Kiley and Beinert 2003; Peña et al. 2013; Liu et al. 2009).

Previously, we demonstrated that *M. gallisepticum* exhibited significant halotolerance and can survive in 1.2 M NaCl solution and grow in 0.5 M NaCl (Gorbachev et al. 2013). The adaptation to hyperosmotic conditions usually involves the transport of low-molecular-weight osmoprotectors, including proline and/or glycine-betaine, via specialized transporters: ProP in *E. coli* (MacMillan and Alexander 1999), BetP in *Corynebacterium glutamicum* (Krämer 2009) and OpuA in *B. subtilis* (Patzlaff et al. 2003). These transporters were not detected in the *M. gallisepticum* genome, suggesting an alternative adaptation mechanism. The genes that were up-regulated under hyperosmotic conditions are enriched in the “aldehyde dehydrogenase [NAD(P)+] activity,” “cellular aldehyde metabolic process,” “acetyl-CoA metabolic process,” “endonuclease activity” and “response to stress” terms (Supplementary Table S5).

Compared with the oxidative and osmotic stresses, the heat stress induced the strongest transcription response. Thus, we studied the kinetics of the heat stress response for 5, 15 and 30 min. The number of heat shock-affected genes increases with the stress duration (Fig. 4A); in total, 538 genes show altered expression for at least one time point. To further classify the heat shock-related expression changes, we clustered the genes into 9 clusters based on the similarity of their expression changes under heat shock (see Methods and Fig. 4B). Most of the genes (for a total of 69%) correspond to continuous expression changes. Among them 232 genes increased expression (clusters 2 and 3) and 136 genes decreased expression (cluster 1). GO enrichment analysis revealed that the translation-associated genes as well as the genes involved in glycolysis were down-regulated (cluster 1), whereas the genes linked with “serine-type endopeptidase activity” and “transposase activity” were quickly up-regulated under heat shock (cluster 2). The genes that were activated later during the heat stress (cluster 3) are involved in “carbohydrate transport” and some other functions. The genes involved in GTP catabolism are quickly inactivated under heat shock (cluster 4; Supplementary Table S7). At the same time, we observed no significant transcriptional changes during the heat stress applied to the cells at the stationary growth phase.

The analysis described above showed that a sufficient proportion of transcription in *M. gallisepticum* corresponds to non-coding regions. Fifty-six (71%) of the non-coding operons overlap with 31 coding operons in the antisense orientation. Most of the antisense operons (47) reside completely within the coding operons. We used the intervals identified above to examine the differential expression of non-coding transcripts. We repeated the analysis described above at the level of intervals, focusing on the 657 expressed intervals that do not overlap with known non-coding genes (rRNA, tRNA and tmRNA). Strikingly, under all stress conditions, the non-coding intervals were significantly more frequently activated than the coding intervals (Fisher exact test, p < 3e-5, 5e-40, 3e-37, 0.02 and 2e-11 for 5, 15 and 30 min of heat shock and for the oxidative and osmotic stresses, respectively).

### Analysis of the protein translation in *M. gallisepticum*

We used LC-MS/MS mass spectrometry to profile the *M. gallisepticum* proteins under the control conditions and after 5, 15 and 30 min of heat shock, and we obtained 278,615 spectra. Using these spectra, we performed an annotation-guided identification of peptides (see Methods), resulting in 6619 peptides from 622 genes. We used these data to calculate the PAI (Ishihama et al. 2008). Our analysis revealed a relatively high protein-to-RNA correlation (pro > 0.55): the PAI values measured under both the control and heat shock conditions correlate with the control levels of mRNA. The protein-to-RNA correlation drops during heat shock because the mRNA abundances change dramatically, whereas the protein levels seem to remain unaffected by such short periods (Fig. 1E).

To identify the peptides that were most likely missed in our annotation, we performed a genome-wide peptide search (see Methods). The procedure resulted in 5634 peptides, most of which (5475) are tryptic peptides that originate from the known proteins. Thirty-three (of the 159 remaining) peptides originated from the ORFs of known genes and correspond to an ORF extension (by usage of the upstream start codon) of seven genes. In two of these seven genes, the extensions are supported by more than one peptide. Most (87) of the remaining 126 peptides are confirmed by only a single spectrum. Grouping these peptides by ORF revealed only four short ORFs supported by more than one peptide. All these ORFs appear to be part of known proteins disrupted by a frame shift that can be explained by sequencing errors.

The results agree with the lack of unannotated CDSs. Consequently, 79 operons with a total length of 115,590 nt (1.3% of mapped reads, excluding tRNA or rRNA reads) with no protein-coding ability are either transcription junk or serve other functions.

To determine the strength of ribosome-binding sites (RBS), we used the RNAduplex program to search for the best (in terms of free energy) RNA duplex that was formed by the 3’-terminal region of 16S rRNA (UUA**CCUCCU**UUCU, homologous to slightly extended Shine-Dalgarno sequence-binding region of *E. coli* (Shine and Dalgarno 1974)) and 100 nt regions upstream of annotated CDSs. Duplexes with a free energy below −8 kcal/mol show pronounced enrichment in the (−25,−1) region (Supplementary Fig. S7). We used the (−100,−76) region as a negative control to estimate the false-discovery rate (FDR), and we found RBSs for 160 ORFs at an FDR of 40%. When the number of RBSs was restricted to 100, the FDR dropped to 30%. The duplex free energy exhibits a weak but significant correlation with the average abundance of both protein and mRNA (rho=−0.22 and −0.27; p < 2e-7 and 1e-10, respectively).

### Transcription regulation

Molecular mechanisms must be responsible for the observed dramatic changes in mRNA abundance under heat and other stresses. In the *Mycoplasma* species, the only TF with a known binding site is a heat-shock repressor, HrcA (GCW_02005), that binds a conserved inverted repeat TTAGCACTC-N_9_-GAGTGCTAA known as CIRCE (controlling inverted repeat of chaperone expression) (Zuber and Schumann 1994; Chang et al. 2008). A genome-wide scan for CIRCE elements that allowed up to two mismatches revealed five CIRCE elements in *M. gallisepticum S6* (Supplementary Table S8). Four of these elements are located upstream of the chaperone gene promoters (*clpB, dnaK, lon* and *dnaJ_2*). The remaining element is located in a pseudogene (*dnaJ* homolog split into three ORFs) promoter region. Logically, the respective genes are up-regulated under heat stress (Fig. 4C). However, their expression fold changes are not significantly higher than those of the other heat-shock up-regulated genes.

We applied the edgeR (Robinson et al. 2010) package to the 5’-ERS read counts to identify TSSs with significant activity changes (BH-corrected p < 0.05 and fold change above two) under heat shock (see Methods). We divided all TSSs into six non-overlapping classes by the mean activity (high or low) and by the change direction: up, not significant and down (Fig. 5A). We applied the MEME suite (Bailey and Elkan 1994) to the sequences that were within 100 nt up- or downstream of CDS-associated TSSs from all six groups but, unfortunately, found no enriched motifs. Even the CIRCE elements discussed above could not be identified, most likely due to their low abundance.

**Figure 5.**
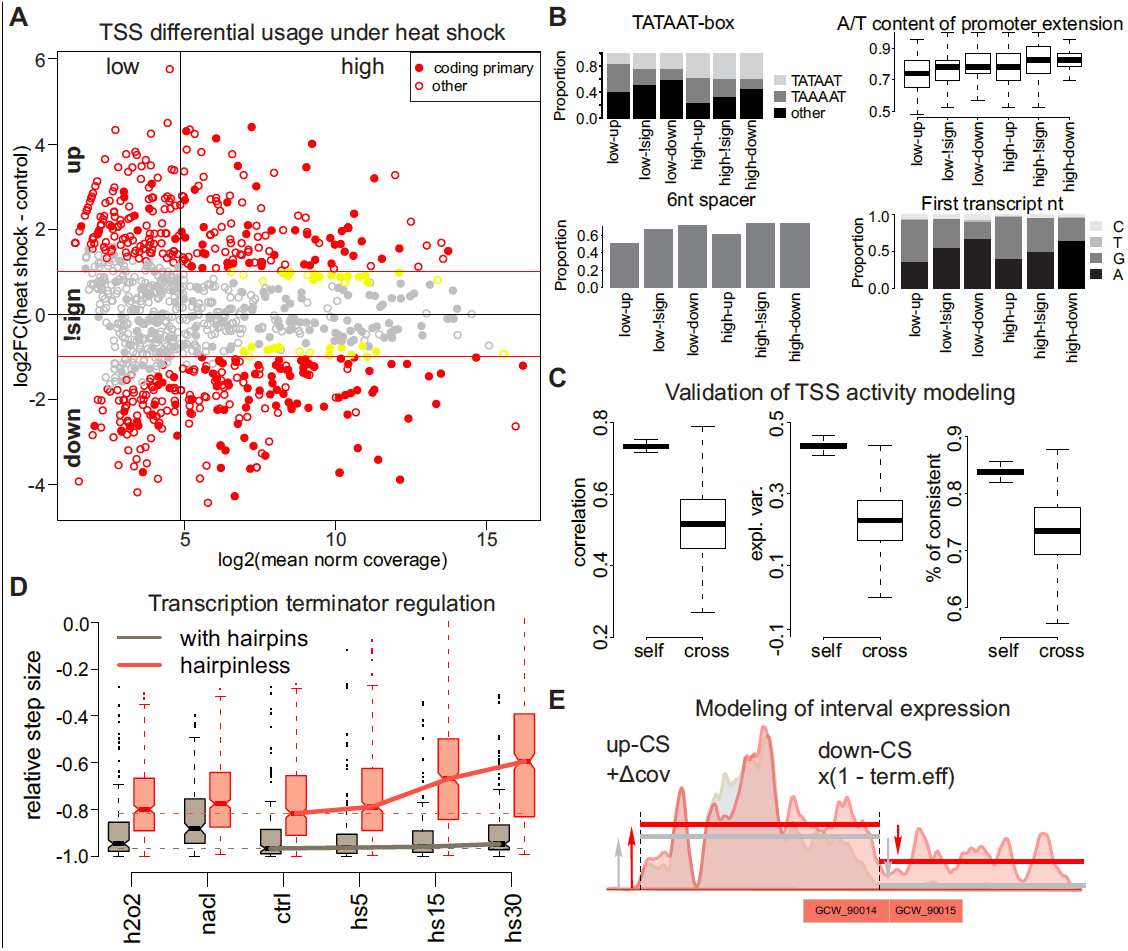
TSS and TT contributions to gene expression regulation. A) Fold change vs. mean activity plot (log scale) for TSSs under heat shock. The TSSs associated with the coding genes are shown with filled circles, the TSSs with significant changes and with |FC|>1 are shown in red, and the TSSs with significant changes but with |FC|<1 are shown in yellow. The TSSs are divided into six groups by the change direction (up, down and not significant (denoted by ‘!sign’)) and by the average activity (low or high; the vertical line represents the median). B) Distributions of TSS properties among the six TSS groups defined in A: type of −10 box, A/T content of the −10 element extension, proportion of promoters with a 6 nt-long spacer between the −10 element and the TSS and frequencies of first transcript nucleotide. C) Pearson correlation, proportion of explained variance and proportion of TSSs with the correct direction of change prediction for the self- and cross-validation of the random forest modeling of the heat-shock fold change. In total, 500 permutations were performed, and in each case, the learning and test sets consisted of 90% and 10% of 495 significant TSSs, respectively. D) Distributions of down-CS efficiency under different conditions and dependence on the presence of hairpins. E) Modeling of the interval expression by additive TSS and multiplicative TT activity. The colored areas represent a smoothed read coverage, the interval borders are shown with vertical lines, and the mean interval coverage is shown with solid lines. The read coverage profiles under the control conditions and heat shock are shown in gray and red, respectively.

Comparing the six TSS groups revealed that the TSSs activated under heat shock differ strikingly from the down-regulated TSSs independently of the mean activity. The heat shock-activated TSSs usually are not associated with CDSs, have a non-canonical spacer (5 or 7 nt) between the −10 element and the TSS, have a lower AT content in the −10 element extension and frequently have a canonical (TAWAAT) −10 element with a preference for TAAAAT (unlike the down-regulated promoters that prefer TATAAT and have a high fraction of noncanonical −10 elements). Additionally, heat-shock-activated transcription is usually initiated from guanosine, whereas the preferred first nucleotide of most down-regulated transcripts is adenosine (Fig. 5B). These observations suggest that a sufficient part of the heat-shock-related expression variation can be explained by these few factors. We used the random forests algorithm (Liaw and Wiener 2002) to predict the log fold change based on the −10 element sequence, the first nucleotide of the transcript, the spacer length and the A/T richness of −10 element extensions (see Methods). The model explained 27% and 14% of the total variance and achieved a Pearson correlation equal to 0.62 and 0.4 in self- and cross-validation, respectively. The correlation reaches 0.52 for cross-validation when only heat-shock-related TSSs are considered (Fig. 5C).

In addition to the transcription initiation that is usually considered a major contributor to transcription regulation, other steps of mRNA synthesis, such as elongation and termination, may also play a role. For example, a change in the efficiency of imperfect TTs might change the expression levels of downstream intervals. In our data, we observed a sufficient drop (by 15%, from −0.72 to −0.56) in efficiency of the hairpinless, but not the hairpin-containing, down-CSs (Fig. 5D). To further investigate the contribution of TTs to transcription regulation, we modeled the read coverage in each interval that resulted from the additive contributions of up-CSs and the multiplicative contribution of down-CSs (see Methods and Fig. 5E). This modeling allowed us to decompose the effects of TSSs and TTs on the expression change of each interval. The results show that for most of the intervals (71% of the 657 expressed intervals that do not overlap with tRNA or rRNA), a single factor (either a TSS or TT) explains more than 80% of the expression variability under heat shock. In general, 87-92% of the transcription variability under heat and other stresses could be explained solely by changes in TSS activity, but there are still many intervals with low expression that are mainly regulated by TTs. In most of the intervals (490), more than 50% of the observed expression variability under heat shock could be attributed to TSSs, whereas in the remaining 167 intervals, the effect of TTs dominates. Of these 167 intervals, 100 change expression significantly under heat shock, and most of them (84%) are activated. The 86 intervals regulated by TTs include 158 CDSs. Of these, 114 change their expression significantly under heat shock. Most of these CDSs (77%) belong to clusters 2 and 3 (Fig. 4B). The genes that are up-regulated under heat shock due to the TT activity change are involved in the process of “carbohydrate transport,” whereas the TSS-regulated genes play a role in transposition, “response to stress” and proteolysis (Supplementary Table S9).

Interestingly, although under most conditions, the hairpin-containing TTs have nearly perfect efficiency (∼ −0.95), their efficiency drops by 8% under osmotic stress (Fig. 5D).

## Discussion

Here, for the first time, we present a genome-wide analysis of the regulation of transcription initiation and termination in *M. gallisepticum* strain *S6*. An analysis of 5’-ERS libraries allowed us to identify 1061 TSSs genome-wide and to decipher their promoter structure. We present a new approach to the analysis of bacterial transcriptomes that is based on the genomic intervals between the two nearest TSSs and/or TTs. This analysis allowed us to separate the genome into 1059 intervals with equal expression levels, to group them into 208 operons and to decompose the effects of transcription initiation and termination on the mRNA concentrations. We have shown a dramatic reorganization of transcription under different biological conditions, such as heat shock or the stationary growth phase. However, other than CIRCE, which was found using *a priori* knowledge, we found no regulatory sequence motifs that could be responsible for the observed regulation. We speculate that other motifs, even if they do exist, would not be found due to their low abundance and thus cannot be responsible for the observed dramatic transcription changes under heat stress.

TAWAAT sequences are avoided in *M. gallisepticum* (they are two-fold less abundant than expected by chance), but their number (3088) is still greater than both: the number of annotated genes and the number of detected TSSs. Only a subset of these sequences is likely to constitute functional promoters. How does the *M. gallisepticum* transcription machinery distinguish the functional promoters from the cryptic ones? We believe that the A/T-richness of the surrounding region, the correct spacer length and the first nucleotide of the transcript play major roles. Increased temperatures may facilitate the recognition of cryptic promoters via the enhanced melting of the DNA in the promoter region. Indeed, heat-shock-activated promoters have a lower AT content, a spacer whose length is not 6 nt and an alteration in the first nucleotide of the transcript (G instead of A). Modeling revealed that these features explain a substantial proportion of the transcription variation under heat shock.

Most of the heat-shock-activated promoters are not associated with CDSs. Because the major fraction of the *M. gallisepticum* genome encodes proteins, the activation of cryptic promoters leads to widespread antisense expression. Interestingly, under the control conditions, the sense and antisense expression exhibit a significant negative Pearson correlation (95% confidence interval is [−0.39, −0.46]), but after 30 min of heat shock, the correlation drops dramatically to [−0.14, −0.27]. Thus, the sense and antisense transcriptions seem to be less coordinated under heat shock, confirming the idea of cryptic promoter activation. We speculate that the down-regulation of promoters during heat stress may be the result of promoter competition for RNA polymerase.

Our analysis of transcription termination reveals two classes of TTs: strong, hairpin-containing TTs and weak, hairpinless terminators. Whereas the former are mostly not regulated during 30 min of heat shock, the efficiency of the hairpinless TTs drops dramatically, apparently contributing to the “promiscuous transcription” described above. A significant portion of the up-regulated CDSs during heat stress can be attributed to regulated TTs. In addition, 39% of the genes from clusters 2 and 3 are regulated predominantly by TTs. As we demonstrated above, a set of the genes coding for carbohydrate uptake proteins is regulated by TTs rather than TSSs. Carbohydrate transport under heat stress may be adaptive, as carbohydrates are a source of ATP for chaperones. We speculate that the gene expression regulation by TTs may represent a novel mechanism of adaptive regulation in genome-reduced bacteria.

Most of the transcripts activated under heat shock do not encode proteins and, thus, seem to be non-functional. This finding suggests that a significant part of the heat-induced transcription changes is non-adaptive. In contrast, CDSs, whose expression follows specific patterns during heat shock or changes under oxidative stresses, are enriched in the relevant biological functions; this finding indicates a tight balance between the transcription noise and regulation.

The HrcA-CIRCE regulatory system is conserved among Mycoplasmas, and this conservation implies its functional importance. However, under heat-stress conditions, CIRCE-dependent promoters behave similarly to numerous CIRCE-less up-regulated promoters, raising questions about the true function of CIRCE. Non-adaptive stress-induced expression changes were previously identified in different bacterial species (Price et al. 2013). We speculate that under heat shock in *M. gallisepticum*, a few adaptive changes, most likely guided by specific transcription factors, occur on the background of a noise-like response.

Our proteomic study shows that the mRNA and protein concentrations correlate well under the control conditions. However, during heat shock, the mRNA, but not protein, levels change dramatically, resulting in a strong decrease in the mRNA-protein correlation. Such a discrepancy might be explained by the different time scales of mRNA and protein turnover (Maier et al. 2011).

One of the greatest weaknesses of current RNA-Seq techniques is their strong dependence of the sequencing efficiency on the sequence and, especially, on the secondary structure of RNA, resulting in a highly variable read coverage depth (Khrameeva and Gelfand 2012). In the current work, we attempted to address this issue by pooling all samples together to minimize the noise during operon prediction and allowing for overdispersion by using a quasi-Poisson distribution. As a side effect of this approach, we have likely lost certain condition-dependent information. The relatively low agreement between the TSSs predicted using up-CSs and those predicted using 5’-ERS might also be explained by a high coverage variability that may lead to both the identification of false up-CSs and the loss of correct TSSs (type I and type II errors, respectively).

In summary, we have taken a step toward understanding the stress-response mechanisms in *M. gallisepticum* and in genome-reduced bacteria in general. Our interval-based approach allowed us to look beyond the operon concept, identify two classes of transcription terminators and decipher their roles in transcription regulation. We believe that the constructed transcription map together with a comprehensive list of TSSs and TTs in *M. gallisepticum* will enhance future research.

## Methods

### Cell culturing

*Mycoplasma gallisepticum S6* was cultivated on a liquid medium containing tryptose (20 g/L), Tris (3 g/L), NaCl (5 g/L), KCl (5 g/L), yeast dialysate (5%), horse serum (10%) and glucose (1%) at pH = 7.4 and 37°C in aerobic conditions and exposed to stress conditions as described previously (Gorbachev et al. 2013).

### RNA extraction

Aliquots of the cell culture were directly lysed in TRIzol LS reagent (Life Technologies) at a 1:3 ratio of culture medium:TRIzol LS (v/v). The lysates were extracted with chloroform, and the aqueous phase was purified with a PureLink RNA Mini Kit (Ambion) to remove tRNA or was used directly to precipitate RNA by the addition of an equal volume of isopropanol.

### Real-time PCR

RNA was treated by DNAse I (Thermo Scientific), and cDNA was synthesized from random hexamer primers by H-minus Mu-MLV reverse transcriptase (Thermo Scientific). Real-time PCR was performed using iQ SYBR Green Supermix (Bio-Rad) and a CFX96™ Real-Time PCR Detection System (Bio-Rad) PCR machine. Quantitative data were normalized to the 23S rRNA transcript as described previously (Gorbachev et al. 2013).

### Preparation of libraries for RNA-Seq

RNA (either total or tRNA-depleted) was fragmented into 200 bp by chemical fragmentation (100 mM ZnSO_4_, 100 mM Tris, pH = 7.0 at 70°C for 15 min). The fragmentation reaction was stopped with 20 mM EDTA (pH = 8.0). The fragmented RNA was treated with T4 polynucleotide kinase (Thermo). Strand-specific double-stranded cDNA libraries for standard RNA-Seq on a SOLiD platform were prepared according to the manufacturer’s protocol using a Total RNA-Seq Kit and a SOLiD RNA Barcoding Kit (Ambion). The quality of the RNA, fragmented RNA and cDNA libraries was assayed with an Agilent 2100 Bioanalyzer system (Agilent).

Amplified ds-cDNA was subjected to a normalization procedure with DSN (double-strand specific nuclease, Evrogen). First, 400-1000 ng of ds-cDNA (12 μL) was mixed with 4 μL of hybridization buffer (200 mM HEPES, 2 M NaCl, pH = 7.5). The procedure was performed in a PCR thermocycler. The samples of ds-cDNA were denatured at 98°C for 2 min and then re-annealed at 68°C for 5 hours. Then, 32 μL of DSN 2X master-buffer (Evrogen) that was prewarmed to 68°C was added. The mixture was incubated at 68°C for 10 min. Subsequently, 0.5 μL of DSN enzyme was added, and the samples were incubated at 68°C for 15 min. The μ reaction was stopped by the addition of 64 μL of 200 mM EDTA (pH = 8.0). Then, an equal μ volume of isopropanol was added, the samples were incubated at −20°C overnight and the cDNA was recovered by centrifugation (20 min at 4°C, 16,000 rcf). Subsequently, the cDNA was amplified and purified using a PureLink PCR Micro Kit (Invitrogen). Then, the procedure was repeated, resulting in two rounds of normalization in total. The normalized cDNA was selected for size by agarose gel electrophoresis (2% agarose, 1xTBE, 4 V/cm). Sample cDNA in the 200-300 bp range was extracted from the agarose blocks using a SOLiD Library Quick Gel Extraction Kit (Life Technologies; E1 buffer from the SOLiD Library Column Purification Kit was used for cDNA elution) and used for downstream preparations according to the standard protocol.

All samples were prepared in two biological replicates with one technical replicate per biological replicate.

### Preparation of libraries for 5′-ERS

To prepare 5’-enriched libraries, at least 20 μg of total RNA was fragmented, end-repaired (as described above) and treated with Terminator exonuclease (Epicentre). This process resulted in the degradation of the non-5’ RNA fragments, whereas the 5’-fragments were protected by the pyrophosphate groups on their 5’-ends. Then, the RNA was precipitated by isopropanol and treated with tobacco acid phosphatase (Epicentre) to remove the pyrophosphate groups. Next, the RNA was precipitated by isopropanol and used for strand-specific ds-cDNA preparation according to the standard protocol for SOLiD libraries. The sample cDNA was normalized in one round as described above and used to prepare SOLiD libraries according to the standard protocol.

### Sequencing

Sequencing was performed on a SOLiD 4 (Life Technologies) platform using SOLiD EZ Bead E80 System Consumables and SOLiD ToP Sequencing Kit, MM50 (Applied Biosystems).

### Protein extraction and 1D electrophoresis

Cells harvested by centrifugation at 10,000 ×*g* at 4°C for 10 min were washed twice in a wash buffer (150 mM NaCl, 50 mM Tris-HCl, 2 mM MgCl_2_, pH = 7.4). The cells were lysed in 20 µL of 1% SDS in 100 mM NH_4_HCO_3_ and incubated in an ultrasonic bath for 15 min followed by centrifugation at 10,000 ×*g* at 4°C for 5 min. The supernatant was extracted, and the protein concentration was determined using a Bicinchoninic Acid Protein Assay Kit (Sigma). Next, 20 µL of 2× Laemmli reagents was added, and the samples were incubated at 95°C for 5 min. Then, 50 µg of protein was loaded onto a polyacrylamide gel (10 × 0.1 cm, 12% polyacrylamide), and electrophoresis was performed as described by Laemmli (Laemmli 1970) (10 mA current). The electrophoresis was stopped when the front dye reached 1.5 cm in the separating gel.

### Trypsinolysis in polyacrylamide gel

The polyacrylamide gel was fixed in a fixation buffer (20% CH_3_OH and 10% CH_3_COOH) for 30 min and washed twice in H_2_O. The gel was cut into 1 × 1 mm pieces, transferred into tubes and treated with 10 mM DTT and 100 mM NH_4_HCO_3_ for 30 min at 56°C. Then, the proteins were alkylated with 55 mM iodoacetamide in 100 mM NH_4_HCO_3_ for 20 min in the dark. Next, water was removed from the gel pieces by the addition of 100% acetonitrile.

The dehydrated samples were treated with a 150 µL of trypsin solution (40 mM NH_4_HCO_3_, 10% acetonitrile, 20 ng/µL Trypsin Gold, mass spectrometry grade; Promega). The samples were incubated for 60 min at 40°C and for 16-18 h at 37°C. Peptides were extracted once by 5% formic acid and twice by 50% acetonitrile with 5% formic acid. The extracts were joined and dried in a vacuum centrifuge at 45°C. The precipitate was diluted in 50 µL of 5% acetonitrile with 1.1% formic acid.

### Chromato-mass spectrometry

The peptides were analyzed using a TripleTOF 5600+ (ABSciex) mass spectrometer with a NanoSpray III ion source and a NanoLC Ultra 2D+ chromatograph (Eksigent). Chromatographic separation was performed in a gradient of acetonitrile in water (5 to 40% of acetonitrile in 120 min) with 0.1% formic acid on 75 × 150 µm columns with a Phenomenex Luna C18 3 µm sorbent and a flow rate of 300 nL/min.

The IDA mode of the mass spectrometer was used to analyze the peptides. Based on the first MS1 spectrum (the mass range for the analysis and subsequent ion selection for MS2 analysis was 300-1250 m/z; the signal accumulation was 250 ms), 50 parent ions with maximum intensity in the current spectrum were chosen for the subsequent MS/MS analysis (the resolution of the quadrupole unit was 0.7 Da, the mass measurement range was 200-1800 m/z, the ion beam focus was optimized to obtain maximal sensitivity, and the signal accumulation was 50 ms for each parent ion). Nitrogen was used for collision dissociation with a fixed average energy of 40 V. The collision energy was linearly increased from 25 to 55 V during the signal accumulation time (50 ms). The parental ions that had already been analyzed were excluded from the analysis for 15 sec.

The mass spectrometry proteomics data have been deposited in the ProteomeXchange Consortium (http://proteomecentral.proteomexchange.org) via the PRIDE partner repository (Vizcaíno et al. 2009) with the dataset identifier PXD000922 and DOI 10.6019/PXD000922.

### Analysis of mass spectrometry data

Raw data files (.wiff file format) were converted to the Mascot generic format (.mgf file format) using AB SCIEX MS Data Converter version 1.3 and were searched using Mascot version 2.2.07 against a database of all proteins (836 amino acids sequences) of *M. gallisepticum S6* (GI:604957178). The Mascot searches were performed with the following parameters: tryptic and semi-tryptic peptides; maximum of one missed cleavage; a peptide charge state limited to 1+, 2+ and 3+; a peptide mass tolerance of 10 ppm; a fragment mass tolerance of 0.5 Da; and variable modifications caused by oxidation (M) and carbamidomethylation (C). Protein scores greater than 6 for protein-trypsin, 17 for protein-semitrypsin, 32 for genome-trypsin and 24 for genome-semitrypsin were assumed to be significant.

The proteogenomic profiling of *M. gallisepticum* was performed using the database of the chromosomal DNA sequence (GenBank) split by 3000 nucleotides with a shift of 1000 nucleotides (493 nucleotides sequences). We used the Protein Abundance Index to evaluate the protein concentrations (as described elsewhere (Ishihama et al. 2008)).

### Read mapping

All reads from DS1 with average quality values below 15 were discarded. Because many reads contain an adapter sequence in the 5’-end, we truncated all reads from both datasets to the first 25 read bases. Then, the reads were mapped to the *M. gallisepticum* strain *S6* genome (GI:604957178) using Bowtie software (Langmead et al. 2009) with the following parameters: bowtie --trim3 23 -f -C -v 3 -y -a --best --strata -S. Each match for the reads that was mapped to multiple positions was treated as an independent read. The results were nearly the same when only the uniquely mapped reads were used.

### Transcription interval identification

To divide the *M. gallisepticum* genome into transcription intervals (intervals with a constant expression level), we combined all RNA-Seq samples from DS2 together and calculated the number of reads that mapped to each position in the genome (read coverage). A read was considered to be mapped to the given position only if its alignment started from that position. Then, we looked for local changes in the read coverage using a sliding window. For each genome position, we calculated the following function:

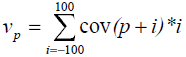

where *cov(j)* is the read coverage at a specific position and *p* is genome position. We considered all local maxima and minima of *v_p_* as possible borders of transcription intervals. Then, we sorted all borders by *v_p_*, and, for each border (starting from borders with highest *v_p_*), we removed all other borders within 25 nt. We used the intervals between the borders to estimate the dependence of the overdispersion parameter on the logarithm of the average coverage using generalized linear models (GLMs) and a quasi-Poisson distribution, *od(cov)*. Then, we applied the following iterative procedure to merge the intervals with similar coverage together:

1. Use GLM with a quasi-Poisson distribution to model the read coverage in each pair of consecutive intervals.
2. Calculate the overdispersion parameter as a maximum of one, the overdispersion from the GLM model calculated in step 1 and the approximation by *od(cov)* calculated for each of two intervals.
3. Find the pair of intervals with the highest p-value (quasi-log likelihood test). Join these intervals if the p-value is lower than 0.05 divided by the genome length.
4. Repeat steps 1-3 until step 3 results in interval merging.

Each remaining border was called a coverage step, either up or down (up-CS and down-CS). For each coverage step and each sample (all union of samples), we calculated its size:

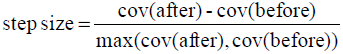

where *cov(after)* and *cov(before)* are the average coverage in interval after and before the step, respectively.

To detect the expressed intervals, we defined the coverage cutoff as the 5% quantile of the average coverage of the CDS-containing intervals. We defined an operon as a set of consecutive expressed intervals.

### TSS identification

To identify TSSs, we used the 5’-ERS data from DS2. Considering each sample separately, we searched for a local maximum in the read coverage (defined as described in the section on transcription interval identification) that was supported by at least 5 reads. Then, we modeled the coverage at each local maximum while considering 5 nt in each direction as background using a GLM with a quasi-binomial distribution and controlling the overdispersion parameter to be not lower than 1. We used a quasi-log likelihood test to identify significant coverage peaks (BH-corrected p-value < 0.05). As a result, 32,148 peaks were detected in at least one (out of four) 5’-ERS sample from dataset 2.

To investigate promoter structure, we focused on the peaks that lie within 300 nt from the nearest start codon and are supported by at least 200 reads. We observed a TATAAT-like sequence surrounded by an AT-rich region in the upstream peak. We built a PWM for the (−32,−3) peak region and then optimized it using the following procedure:

1. For each peak, find the best PWM match in the (−34,−1) region.
2. Order the peaks by match weight and take the top 60%.
3. Rebuild the PWM using the matches selected in step 2.
4. Repeat steps 1-3 until convergence is achieved.

The optimized PWM matrix is shown in Supplementary Table S9.

Then, we searched for the best match of the resulting PWM in all peaks. To this end, we separated the PWM into two parts: PWM1 corresponded to the TATAAT-like region, and PWM2 corresponded to the surrounding region. We first looked for positions with the best PWM1 match; if there was more than one position that had a maximum score, we selected the one with the highest PWM2 score. The results (Supplementary Fig. S5) show that the positions that correspond to a 5-7 nt spacer between the TATAAT box and the TSS are preferred. Thus, we used these positions to predict the best promoter sequence for each peak. To evaluate the background distribution of the weight of both PWM parts, we scanned the whole genome. For every three consecutive positions, we chose the best PWM match. We divided the whole range of PWM1 and PWM2 scores into 20 bins that resulted in a 2D (20 × 20) PWM score distribution. In each bin of PWM1, we set a specific cutoff for the PWM2 score to achieve an FDR below 10% (Supplementary Fig. S8), resulting in the identification of 1061 TSSs.

To identify TSSs that have a TRTGN extension or −35 box, we searched for the corresponding sequences (TRTG and TTGACA with no more than two mismatches) in the regions upstream of the TATAAT box. We compared the numbers of matches in the expected regions (−5 and −23 from TATAAT) and in the (−86,−36) region.

CIRCE elements were annotated by a genome-wide search for the TTAGCACTC-N_9_-GAGTGCTAA sequence that allowed for up to two mismatches (Supplementary Table S8).

### Hairpin prediction

We used the RNIE (Gardner et al. 2011) program in sensitive mode to predict hairpins in the *M. gallisepticum* genome.

### Differential expression analysis

The read counts and reads per kilobase of transcript per million mapped reads (RPKM) for the CDSs and intervals were calculated using SAJR software (Mazin et al. 2013). Each location of the reads that mapped to multiple locations was treated independently and in the same manner as the locations of uniquely mapped reads. For most of the genes, the results did not change when only the uniquely mapped reads were considered. Only the expressed intervals that did not overlap with any known non-coding (mtRNA, tRNA and rRNA) genes were used. Library sizes were calculated as the sums of the coverage of all CDSs (or intervals). All library sizes (except for the two stationary samples) were adjusted by RLE normalization using the package edgeR (Robinson et al. 2010). Then, we applied SAJR to detect differentially expressed genes while considering the number of reads mapped to a given CDS (or interval) as the result of the binomial trials with the number of trials equal to the library size. The CDSs (and intervals) with a Benjamini Hochberg (BH)-corrected (Benjamini and Hochberg 2009) p-value below 0.05 and with a fold change above two were considered to be significantly differentially expressed.

A similar procedure, with two differences, was used to analyze TSS activity under heat shock: first, for each TSS, only the reads that mapped exactly to a transcription initiation site were used; second, we used the exact test from the edgeR package to assess the significance of the expression change.

### Heat-shock gene clustering

Genes whose expression changed significantly after at least one heat-shock period were clustered using hierarchical clustering with complete linkage and using one minus the Pearson correlation between the expression levels (control and heat-shock samples) as a distance measure. The expression levels (RPKMs) were averaged between the replicates and were logarithmized. The obtained dendrogram was divided to form nine clusters. The resulting clusters were reordered by size. For visualization, RPKMs were z-score transformed. The boxplots in Fig. 4B represent the distribution of z-scores in the given condition for all genes for a particular cluster.

### Gene Ontology (GO) enrichment analysis

GO annotation was conducted using the blast2GO program (Conesa et al. 2005). Enrichment analysis was performed using the goseq package (Young et al. 2010), and all annotated CDSs were used as the background. Only the terms with more than two genes were considered. All terms with a BH-corrected p-value above 0.2 were considered significant.

### Modeling of TSS heat-shock response

We modeled the log fold change of the heat-shock-related TSS activity change using the randomForest package (Liaw and Wiener 2002). The sequence of the −10 box (the sequences that met less than 10 times were considered a single class), the first nucleotide of the transcript and the spacer between the −10 box and the TSS were used as categorical predictors, and the AT contents within 20 nt upstream and 3 nt downstream of the Pribnow box were used as continuous variables. We fitted the model 500 times to 90% randomly chosen TSSs and used the remaining 10% for cross-validation.

### Modeling of the interval expression

We modeled the average coverage in each expressed interval under each condition (replicates were merged together, and the coverage was scaled by the sum of adjusted using RLE normalization (by intervals; see above) library sizes using the following model:

1. The coverage (cov) before the start of each operon, was set to 0
2. At the i^th^ up-CS, the coverage is updated by cov = cov + effect(up-CS_i_), where effect(up-CS_i_) cannot be negative
3. At the i^th^ down-CS, the coverage is updated by cov = cov*effect(down-CS_i_), where effect(down-CS_i_) is between 0 and 1

Such models perfectly explain (with a few negligible exceptions) the interval coverage under each condition. To decompose the effects of TSSs (effect(up-CS)) and TTs (effect(down-CS)) on the expression changes after 30 min of shock, we built two models – one for the control conditions and one for the heat shock. Then, we constructed an intermediate model that utilizes the TSS effects from the model built for the heat shock sample and the TT effects from the control model and used it to model the interval coverage. We considered the difference between the control condition and the results of the coverage prediction by the intermediate model to be TSS-related effects, and we considered the difference between the coverage prediction by the intermediate model and the interval coverage under heat shock to be TT-related effects. The feature with the greatest effect was considered the major driver of the expression change for a given interval. The results changed only moderately when the intermediate model was constructed in the opposite way (using the TSS effects from the control model and the TT effects from the model built for heat-shock samples).

## Data access

*M. gallisepticum S6* genome sequence and annotation were deposited in NCBI GenBank under GenBank id CP006916.2. Transcriptomics data was uploaded to NCBI SRA database under project id PRJNA243934. Proteomics data was uploaded to ProteomeXchange Consortium database via the PRIDE partner repository under dataset id PXD000922 and DOI 10.6019/PXD000922.

